# Inhibition of cell fate repressors secures the differentiation of the touch receptor neurons of *Caenorhabditis elegans*

**DOI:** 10.1101/320978

**Authors:** Chaogu Zheng, Felix Qiaochu Jin, Brian Loeber Trippe, Ji Wu, Martin Chalfie

## Abstract

Terminal differentiation generates the specialized features and functions that allow postmitotic cells to acquire their distinguishing characteristics. This process is thought to be controlled by transcription factors called “terminal selectors” that directly activate a set of downstream effector genes. In *Caenorhabditis elegans* the differentiation of both the mechanosensory touch receptor neurons (TRNs) and the multidendritic nociceptor FLP neurons utilize the terminal selectors UNC-86 and MEC-3. The FLP neurons fail to activate TRN genes, however, because a complex of two transcriptional repressors (EGL-44/EGL-46) prevents their expression. Here we show that the ZEB family transcriptional factor ZAG-1 promotes TRN differentiation not by activating TRN genes but by preventing the expression of EGL-44/EGL-46. Since EGL-44/EGL-46 also inhibits the production of ZAG-1, these proteins form a bistable, negative feedback loop that regulates the choice between the two neuronal fates.

**Summary statement:** Transcriptional repressors regulate binary fate choices through reciprocal inhibition during terminal neuronal differentiation. Specifically, ZEB family transcription factor safeguards fate specification of touch receptor neuron by inhibiting TEA domain-containing repressor.

## Introduction

The terminally differentiated state or cell fate of neurons distinguishes them from other neurons through specialized features and functions and the expression of effector genes, termed as “terminal differentiation genes” (Hobert, 2011). Although individual effector genes may not be expressed in a cell type-specific manner, the collective expression of a battery of terminal differentiation genes serves as a signature for the cell fate. For example, the cell fate of mammalian photoreceptors is defined by the expression of over 600 genes, including those encoding proteins directly involved in light detection (Blackshaw et al., 2001; Hsiau et al., 2007). The expression of the terminal differentiation genes is often activated by transcription factors called “terminal selectors” (Garcia-Bellido, 1975; Hobert, 2011) that act alone or in combination and bind to common *cis*-regulatory elements in the terminal differentiation gene.

Terminal selectors or transcriptional activators are not, however, sufficient to specify neuronal fates. Neuron type-specific transcriptional repressors sculpt the expression profiles of terminal differentiation genes by repressing effector genes that do not contribute to a given fate (Kerk et al., 2017; Mitani et al., 1993; Pflugrad et al., 1997; Vallstedt et al., 2001; William et al., 2003). Such repressors appear to be particularly important since the same terminal selectors are often expressed in several distinct types of neurons and promote their differentiation into distinct fates (Hobert, 2016). For example, although UNC-3 is the terminal selector for all cholinergic motor neurons in the *C. elegans* ventral nerve cord, combinations of class-specific repressors control the subtype identity of those motor neurons by antagonizing the activity of UNC-3 on selective effector genes (Kerk et al., 2017). Because repressors restrict selector activity, their expression needs to be tightly regulated to ensure the proper differentiation of a particular cell fate.

To understand the mechanism that controls the expression of repressors, we have studied the choice of cell fate that generates either the touch receptor neurons (TRN) or the FLP neurons in *C. elegans*. The six mechanosensory TRNs detect gentle mechanical stimuli along the body (Chalfie and Sulston, 1981), whereas the two FLP neurons are multidendritic nociceptors that sense harsh touch, noxious temperature, and humidity (Chatzigeorgiou and Schafer, 2011; Chatzigeorgiou et al., 2010; Kaplan and Horvitz, 1993; Russell et al., 2014).

The heterodimer of the POU homeodomain transcription factor UNC-86 and the LIM homeodomain transcription factor MEC-3 acts as a terminal selector that promotes TRN cell fate (Xue et al., 1993). Specifically, the heterodimer activates a set of TRN terminal differentiation genes (the mechanosensory channel genes *mec-4* and *mec-10*, the tubulin genes *mec-7* and *mec-12*, the tubulin acetyltransferase gene *mec-17*, and others) by binding to conserved regulatory elements in their proximal promoters (Duggan et al., 1998; Zhang et al., 2002).

MEC-3 and UNC-86 are expressed in and needed for the differentiation of the FLP neurons and the two postembryonic, multidendritic PVD neurons. The expression of *unc-86* and *mec-*3 in these cells does not lead to the normal expression of many TRN terminal differentiation genes, resulting in cells whose fate is very different from the TRNs (Finney and Ruvkun, 1990; Way and Chalfie, 1988). Our previous work suggested that the TEA domain transcription factor EGL-44 and the zinc-finger protein EGL-46, which form a complex abbreviated as EGL-44/EGL-46, in FLP neurons prevent these cells from acquiring the TRN fate (Mitani et al., 1993; Wu et al., 2001). Loss of these genes in the FLP cells turns them into TRN-like cells; and misexpression of these genes in the TRNs prevent them from taking on TRN characteristics. Thus, repression by EGL-44 and EGL-46 in FLP neurons distinguished them from the TRNs.

In this paper, we extend this model of TRN and FLP neuronal determination by identifying another cell fate regulator, the Zinc finger homeodomain transcription factor ZAG-1, which is expressed in TRNs but not FLP (or PVD) neurons. ZAG-1 promotes the TRN fate not by directly activating the TRN terminal differentiation genes but by preventing *egl-44* and *egl-46* expression in the TRNs. In FLP neurons, the EGL-44/EGL-46 complex simultaneously represses the TRN genes, activates FLP genes, and represses *zag-1*. The mutual inhibition by EGL-44/EGL-46 and ZAG-1 establishes a bistable switch between TRN and FLP fates. Our work suggests that UNC-86/MEC-3 serves as a ground-state selector resulting in a common state in both TRNs and FLP neurons, and that individual fates are subsequently controlled by this bistable switch. We hypothesize that ZAG-1, particularly, regulates the choice of cell fate by inhibiting fate repressors to safeguard the activation of terminal differentiation genes.

## Results

### An RNAi screen for transcription factors affecting TRN cell fate

The six *C. elegans* TRNs include two pairs of embryonically-derived and bilaterally symmetric neurons (the anterior ALML and ALMR and posterior PLML and PLMR), and the unpaired, postembryonically-derived AVM and PVM neurons (Chalfie and Sulston, 1981; Sulston and Horvitz, 1977). To search systematically for transcription factors specifying TRN fate, we knocked down the expression of transcription factor genes using RNAi and looked for animals that no longer expressed *mec-17p::RFP*, a TRN marker (see Materials and Methods for details). Using RNAi against *unc-86* as a positive control, we tested various genetic backgrounds that were previously found to enhance the effects of RNA interference and found that *eri-1*; *lin-15B* mutants had the highest penetrance for the loss of RFP expression. About 80% of these animals did not express RFP in either of the two ALM neurons when treated with *unc-86* RNAi (Figure 1A and B). The posterior TRNs (PLM neurons) were less affected by the RNAi treatment (Figure 1B), as seen previously (Calixto et al., 2010). Therefore, we focused on the disappearance of *mec-17p::RFP* expression in the ALM neurons in the screen.

**Figure 1.**
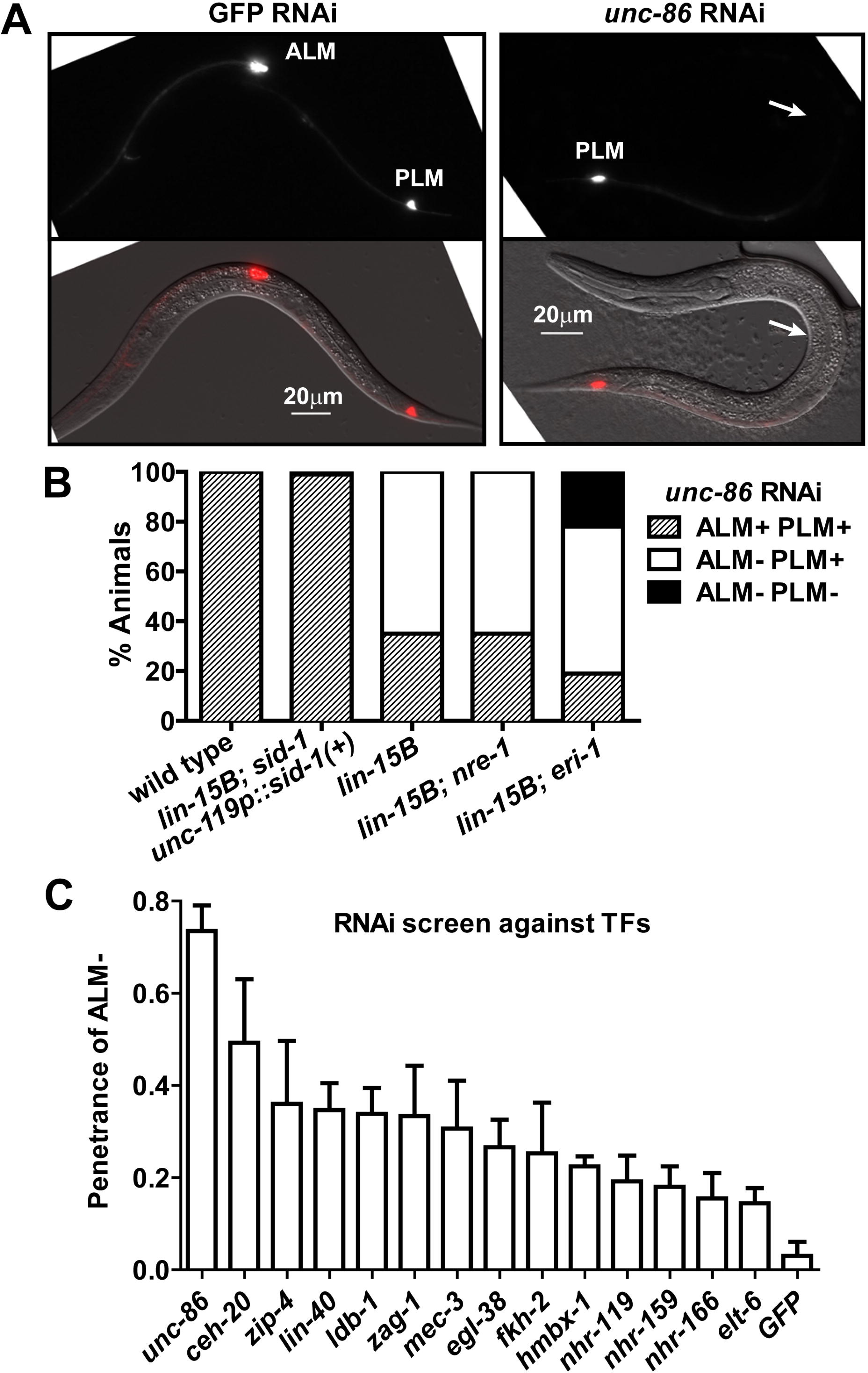
RNAi screen identifies positive regulators of TRN fate. (A) TU4429, *eri-1(mg366); lin-15B(n744); uIs134[mec-17p::RFP]* animals treated with RNAi against *unc-86* or GFP. (B) Percentage of animals that showed RFP or GFP expression in at least one ALM and one PLM (ALM+PLM+), in no ALM but at least one PLM (ALM-PLM+), and in no ALM and no PLM (ALM-PLM-) neurons, respectively. Strains tested for the efficiency of RNAi are TU4429, TU4396, *nre-1(hd20) lin-15B(hd126); uIs134[mec-17p::RFP]*, TU3595, *sid-1(pk3321) him-5(e1490); lin-15B(n744); uIs72[mec-18::GFP]*, and TU4301, *lin-15B(n744); uIs115[mec-17p::RFP]*. (C) The positive RNAi clones identified from the screen. *p* > 0.05 for all the positives using the data from all five rounds of screen.

Among the 443 bacterial clones expressing dsRNA against 392 transcription factors and associated proteins (Table S1), we identified 14 genes that were required for the expression of TRN markers (Figure 1). Four of these genes (*unc-86*, *mec-3*, *ldb-1*, and *ceh-20*) were known to affect the expression of TRN terminal differentiation genes (Cassata et al., 2000; Way and Chalfie, 1988; Zheng et al., 2015b). We examined null mutants for all the remaining ten genes but could only confirm the loss of *mec-17p::RFP* expression in *zag-1* mutants.

We were surprised that RNAi against nine genes affected TRN expression through six rounds of testing but loss-of-function mutations in these genes did not. These false positive results are unlikely to result from the specific genetic background of the RNAi strain, since mutation of several of the genes (*zip-4*, *hmbx-1*, and *nhr-119*) in *eri-1; lin-15B* animals did not affect TRN fate. Activating the RNAi pathway non-specifically by RNAi against GFP in those triple mutants (e.g. *zip-1; eri-1; lin-15B* mutants) did not cause the loss of *mec-17p::RFP* expression either. The discrepancy between the RNAi and mutant phenotypes has been seen in other systems (Kok et al., 2015; Poole et al., 2011) and may be due to mistargeting of the dsRNAs or genetic compensation in mutants (see Discussion for details).

### ZAG-1 is required for the expression of TRN fate markers but acts independently of UNC-86/MEC-3

*zag-1* encodes the sole *C. elegans* homolog of ZEB transcription factors, all of which contain a homeodomain flanked by clusters of C2H2-type zinc fingers. Human ZEB transcription factors induce the epithelial to mesenchymal transition (EMT) and are essential for normal embryonic development; mutations in ZEB genes cause defects in neural crest development and are linked to malignant tumor progression (Vandewalle et al., 2009). *C. elegans zag-1* regulates the differentiation and axonal guidance of several types of neurons, including the command interneurons, GABAergic motor neurons, dopaminergic sensory neurons ADE and PDE, the posterior interneuron PVQ, and the pharyngeal neuron M4 (Clark and Chiu, 2003; Ramakrishnan and Okkema, 2014; Wacker et al., 2003).

Previous research on the TRNs showed that a hypomorphic *zag-1* allele (*zd86*) caused defects in ALM migration and axonal growth (Clark and Chiu, 2003) and made PVM adopt the shape of the multidendritic nociceptor neuron PVD (Smith et al., 2013). Using a *zag-1* null mutation (*hd16*), we found that the complete loss of *zag-1* led to larval arrest in the L1 stage (before AVM and PVM arise) and failure to express several TRN fate markers in the ALM and PLM neurons (Figure 2A-D). The *zag-1* null allele *hd16* deletes part of the first exon and part of the first intron (Wacker et al., 2003), whereas *zd86* and two other viable alleles (*zd85* and *rh315*) all cause premature stops in the fifth exon, leaving the first two zinc fingers and the homeodomain intact (Figure 2B). Thus, ZAG-1 is important for general TRN fate specification.

**Figure 2.**
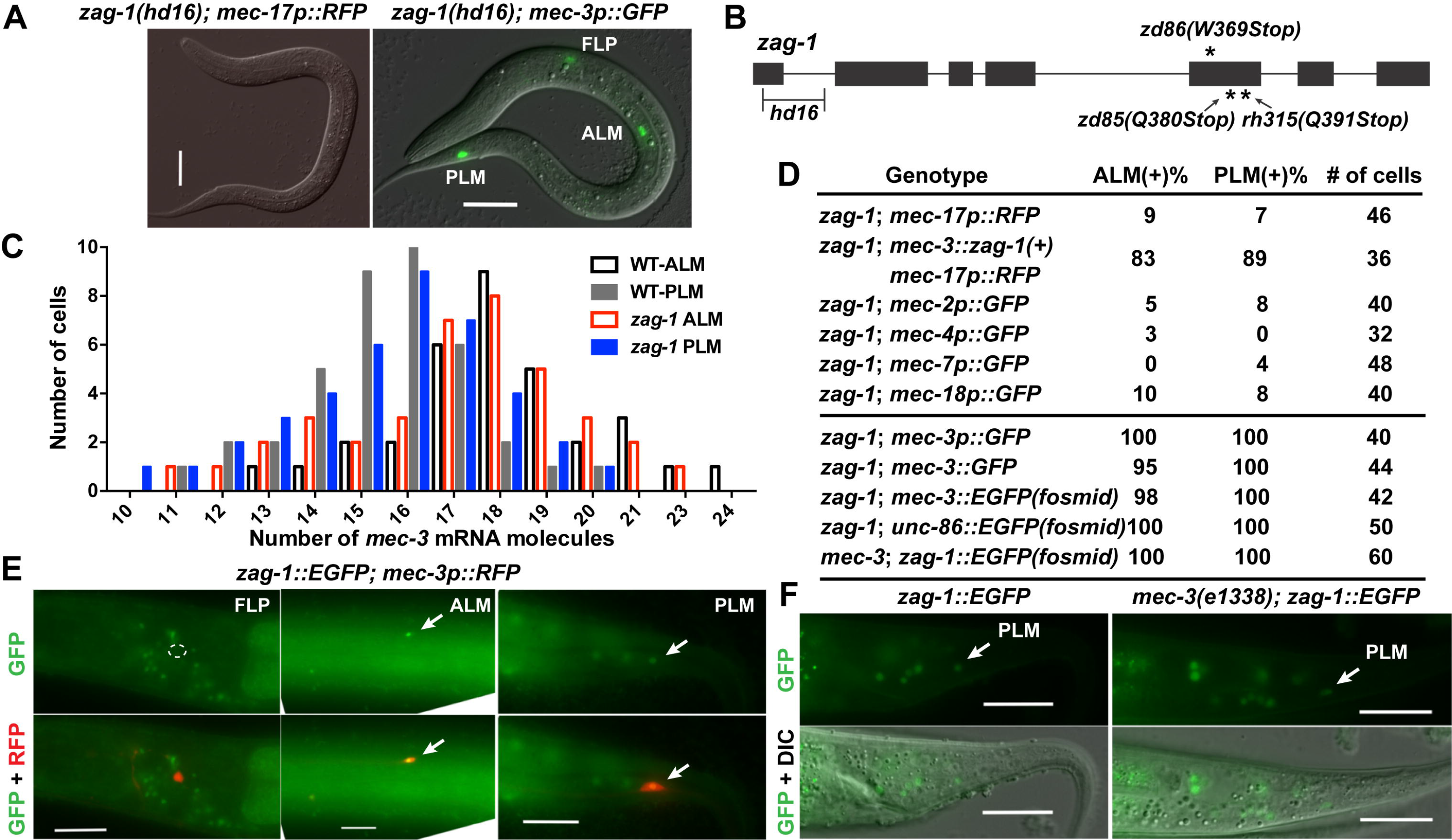
*zag-1* is required for the expression of TRN markers independently of *mec-3*. (A) The expression of *mec-17p::RFP* and *mec-3p::GFP* reporters in *zag-1(hd16)* L1 larvae. (B) The structure of *zag-1* gene and the positions of *zag-1* mutations. (C) The number of *mec-3* transcripts in TRNs from wild-type and *zag-1* animals from smFISH experiments. (D) The penetrance for the expression of the TRN fate markers and *mec-3*, *unc-86*, and *zag-1* reporters in ALM and PLM cells of *zag-1(hd16)* animals at L1 stage. Reporters for those well-characterized TRN effector genes are expressed in 100% ALM and PLM cells in the wild-type animals, and their expression signals the proper differentiation of the TRN fate. We expect most of the terminal differentiation genes associated with the general TRN fate to act similarly to those selected fate markers. (E) The expression of fosmid-based reporter *zag-1::EGFP* in TRNs but not FLPs. (F) *zag-1:EGFP* expression in *mec-3* mutants. Scale bars = 20 m.

We next tested the genetic interaction between *zag-1* and *mec-3* and found that the *zag-1 (hd16)* null allele did not affect the expression of the *mec-3* using a transcriptional reporter (Figure 2A), quantitative mRNA measurements with single molecule fluorescent *in situ* hybridization (smFISH; Figure 2C), or MEC-3::GFP translational fusions (Figure 2D). In addition, expression of *zag-1(+)* from the *mec-3* promoter restored the expression of the TRN fate marker *mec-17p::RFP* in *zag-1* mutants, suggesting that ZAG-1 induces the expression of TRN terminal differentiation genes cell-autonomously. Since *mec-3* expression is completely dependent on *unc-86*, we expected and found that the expression of *unc-86* was not changed in *zag-1* mutants.

Using a fosmid-based GFP translational fusion, we found that *zag-1* was expressed in the six TRNs but not in the FLP and PVD neurons (Figure 2E), and the expression of *zag-1* in TRNs was not affected by mutations in *mec-3* (Figure 2F). Therefore, *zag-1* and *mec-3* are transcriptionally independent of each other. We also failed to find any physical interaction between ZAG-1 and MEC-3 in yeast two-hybrid assays (Figure S1). Together, our results suggest that ZAG-1 promotes TRN fate independently of UNC-86 and MEC-3; the expression of the three transcription factors only overlap in the TRNs and thus form a unique combinatorial code for TRN fate.

Smith *et al*. (2013) found that another conserved transcription factor AHR-1 (aryl hydrocarbon receptor) controls the differentiation of AVM; in *ahr-1* null mutants, AVM cells adopted a PVD-like multidendritic shape. They hypothesized that AHR-1 and ZAG-1 function in parallel to specify TRN morphology in the AVM and PVM, respectively. We found that *ahr-1* is expressed in all six TRNs but is only required for the expression of TRN markers in AVM neurons, suggesting that AHR-1 plays a subtype-specific role in TRN fate specification (Figure S2A-C). In contrast, ZAG-1 is required for TRN fate adoption in general.

### ZAG-1 promotes TRN fate by suppressing TRN fate inhibitors EGL-44/EGL-46

The TEA domain transcription factor EGL-44 and the Zn-finger protein EGL-46 repress TRN fate in the FLP neurons (Wu et al., 2001). These proteins appeared to work together to regulate gene expression and physically interacted in yeast two-hybrid assays (Figure S3). Both genes were normally expressed in the FLP neurons but not the TRNs, and the expression of *egl-46* was dependent on *egl-44* (Wu et al., 2001) (Figure 3A and B). Furthermore, *egl-44* and *egl-46* mutations caused the ectopic expression of *mec-17p::GFP* and other TRN reporters in FLP neurons (Wu et al., 2001); Figure 3A and C; Table S2).

**Figure 3.**
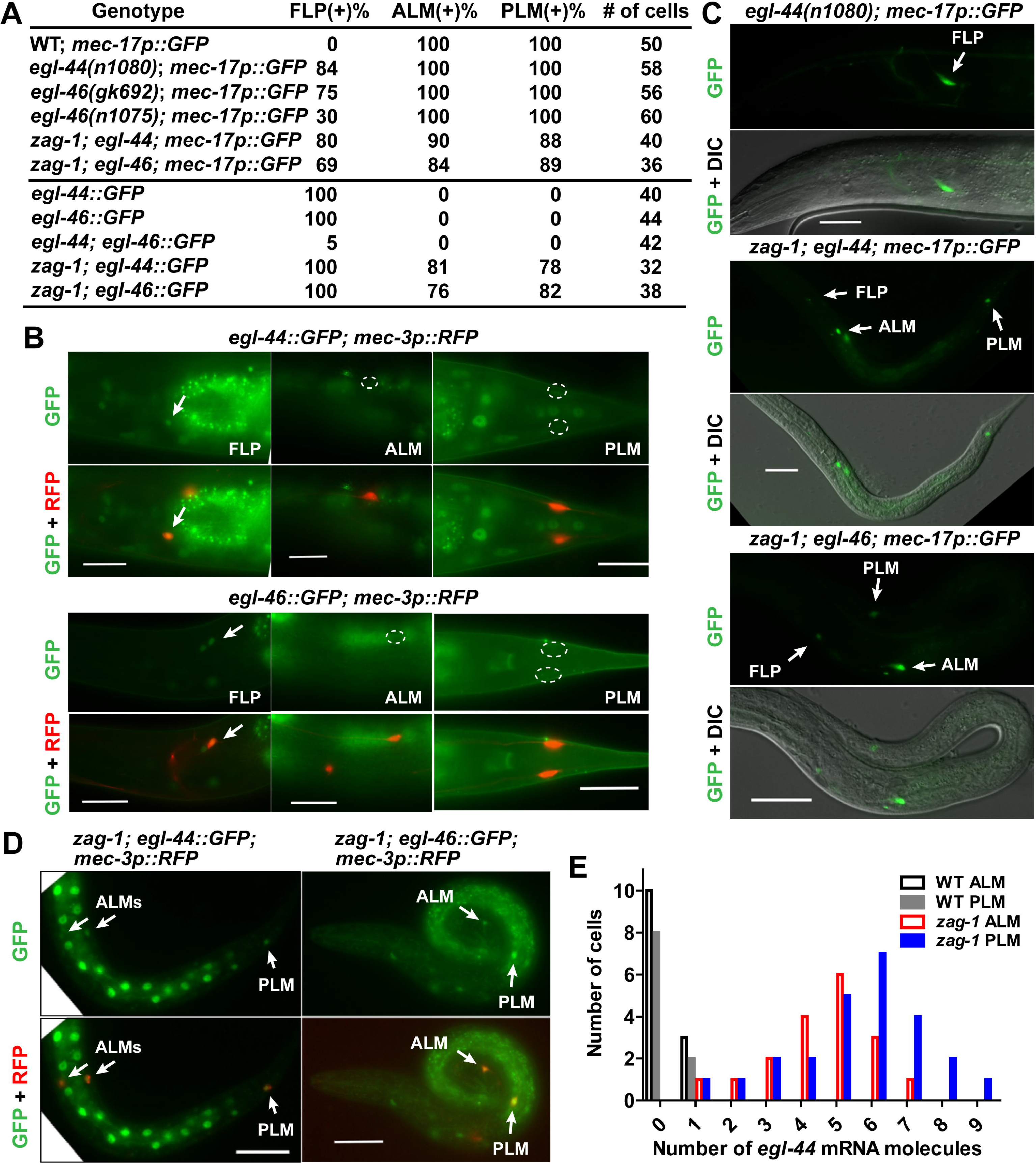
ZAG-1 promotes TRN fate by repressing *egl-44* and *egl-46*. (A) The expression of the TRN marker *mec-17p::GFP* in FLP neurons of *egl-44* animals and in TRNs of *egl-44*; *zag-1* and *zag-1; egl-46* animals. (B) Penetrance of the expression of various reporters. (C-D) The expression of *egl-44::GFP* and *egl-46::GFP* reporters in FLPs but not TRNs of wild-type animals and in TRNs of *zag-1* animals. (E) The number of *egl-44* mRNA molecules in TRNs of wild-type and *zag-1* animals.

Since *zag-1* was selectively expressed in TRNs but not FLP neurons, we tested if ZAG-1 promoted the TRN fate by preventing the activation of EGL-44/EGL-46 in these cells. Consistent with this hypothesis, GFP reporters for both *egl-44* and *egl-46* were ectopically expressed in the TRNs in *zag-1* mutants (Figure 3D). In addition, *egl-44* mRNA, as measured by smFISH, was increased in *zag-1* TRNs, an indication that ZAG-1 transcriptionally repressed *egl-44* (Figure 3E). Importantly, *egl-44* and *egl-46* are epistatic to *zag-1*, since mutations in them restored the expression of TRN fate markers in *zag-1*-deficient animals (Figure 3C and Table S2). Thus, the TRN cell fate did not require ZAG-1 in the absence of EGL-44/EGL-46. Instead, ZAG-1 promoted the TRN fate through a double inhibition mechanism by preventing the expression of the TRN fate repressors *egl-44* and *egl-46*.

We also found that loss of ZAG-1 affected general neurite growth and guidance in TRNs, since *egl-44; zag-1* and *zag-1*; *egl-46* mutants showed shortened and misguided TRN neurites. For example, ALM neurites and the PLM anterior neurites were absent or severely shortened in about 50% and 70% (N = 40) of those double mutants, respectively (Figure S4). Our results using the *zag-1* null allele revealed its essential role in regulating neurite development, which extends previous observation that hypomorphic *zag-1* alleles caused axonal guidance defects in multiple neurons (Clark and Chiu, 2003; Wacker et al., 2003). Because these outgrowth defects are not seen in *mec-3* mutants, ZAG-1 has a second, independent function in TRNs in addition to inhibiting *egl-44* and *egl-46*.

### Misexpression of ZAG-1 converts FLP neurons into TRN-like cells

To address whether *zag-1* could affect FLP differentiation, we misexpressed *zag-1* in FLP neurons using the *mec-3* promoter. This misexpression led to the expression of *mec-17p::GFP* and other TRN markers (Figure 4A and Table S2) and diminished expression of *egl-44* and *egl-46* (Figure 4A and B) in FLP neurons. These data suggest that misexpressed ZAG-1 converts FLP neurons to a TRN-like fate by inhibiting *egl-44* and *egl-46*. The expression of *mec-3p::zag-1* transgene also activated TRN markers in PVD neurons (Figure 4A). Thus, ZAG-1 not only prevents PVM neurons from taking on the PVD fate (Smith et al., 2013) but can also turn PVD neurons into TRN-like cells. Since *egl-44* was not expressed in PVD neurons, the misexpressed ZAG-1 presumably directs this conversion of cell fate by inhibiting some unidentified factor(s) that normally prevents the acquisition of the TRN fate by PVD neurons.

**Figure 4.**
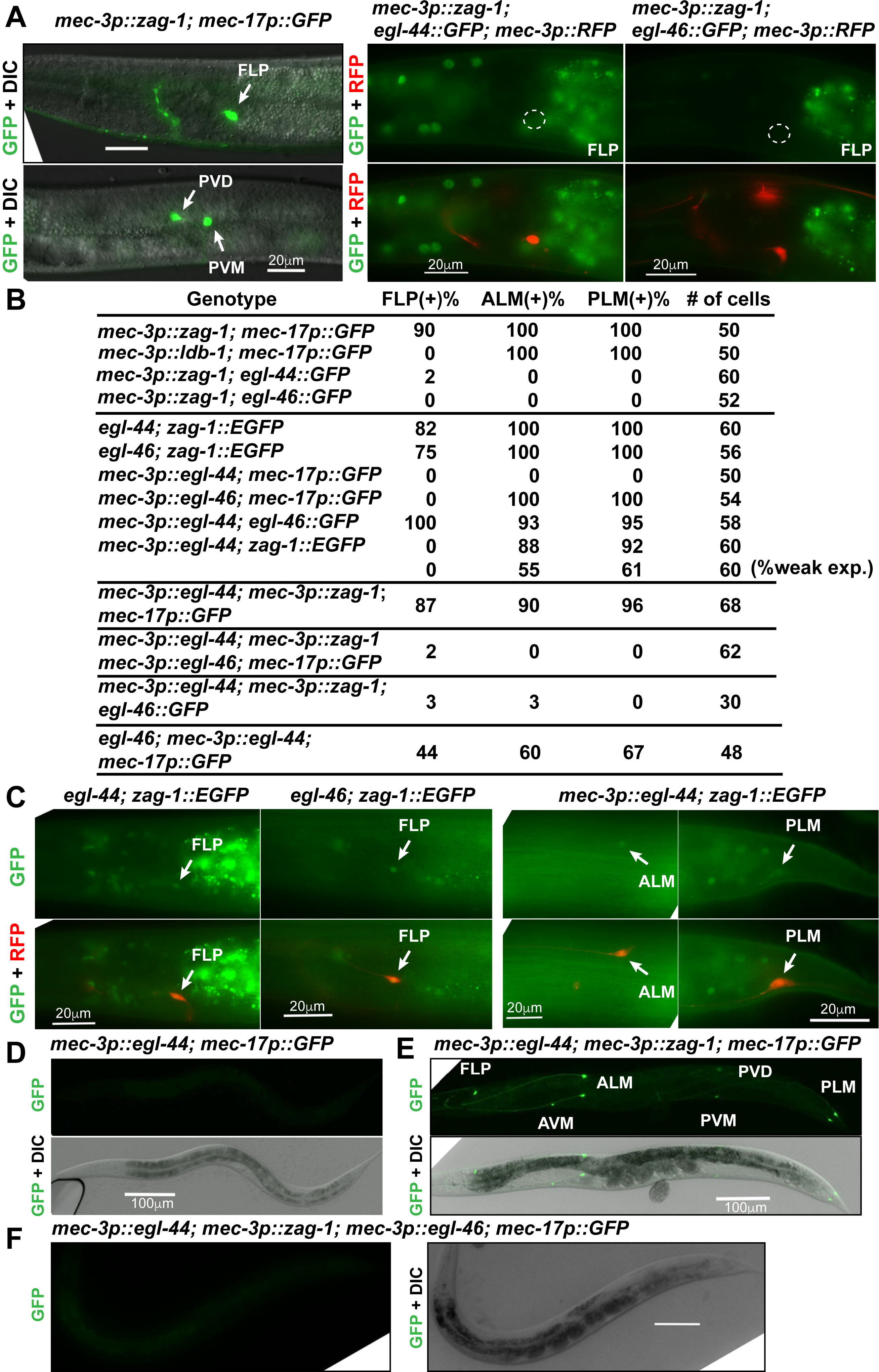
Mutual inhibition between *zag-1* and *egl-44* regulates TRN fate decision. (A) The activation of the TRN marker *mec-17p::GFP* and the loss of the expression of *egl-44* and *egl-46* reporters in FLP neurons of animals carrying *mec-3p::zag-1* transgene. (B) The percentage of cells expressing the indicated markers in various strains. (C) The expression of *zag-1::EGFP* in FLP neurons of *egl-44* and *egl-46* mutants, and weak expression of *zag-1* reporter in animals misexpressing *egl-44* from the *mec-3* promoter. (D-F) The expression of *mec-17p::GFP* in animals carrying various transgenes.

We also expressed *ahr-1* from the *mec-3* promoter and found that misexpressed AHR-1 could activate TRN fate markers in PVD but not FLP neurons (Figure S2D-E). Overexpression of AHR-1 in TRNs also caused morphological defects, such as the growth of an ectopic ALM posterior neurite (Figure S2D), which does not occur when ZAG-1 was overexpressed. These results further support the hypothesis that ZAG-1 and AHR-1 have different functions.

### EGL-44/EGL-46 prevents *zag-1* expression in FLP neurons

Given the mutually exclusive patterns of *zag-1* and *egl-44/egl-46* expression in FLP neurons and TRNs, we next tested whether EGL-44/EGL-46 regulated *zag-1* expression. Mutations in *egl-44* and *egl-46* resulted in ectopic expression of a *zag-1::EGFP* reporter in FLP neurons. Misexpression of *egl-44* from the *mec-3* promoter is sufficient to suppress the TRN fate, because it activates the endogenous *egl-46*, a cofactor that is required for EGL-44 functions (Wu et al., 2001) (Figure 4B and D). We found that EGL-44 misexpression reduced, but did not completely eliminate, the expression of *zag-1* in TRNs (Figure 4B and C). Therefore, the positive TRN fate regulator ZAG-1 and the negative regulator EGL-44/EGL-46 reciprocally inhibit each other’s expression.

ZAG-1 also inhibited endogenous *egl-46* expression. When EGL-44 and ZAG-1 were simultaneously misexpressed from the heterologous *mec-3* promoter, the presence of both proteins led to the activation of TRN markers in all *mec-3*-expressing neurons (TRNs, FLP, and PVD; Figure 4E), because ZAG-1-mediated inhibition of endogenous *egl-46* blocked the effects of EGL-44. Expression of an *egl-46*::GFP reporter was reduced by ZAG-1 despite the presence of misexpressed EGL-44 (Figure 4B), suggesting that ZAG-1 can repress *egl-46* transcription both through *egl-44* and independently of *egl-44*. To rule out the possibility of direct protein-protein interaction and interference, we misexpressed EGL-44, EGL-46, and ZAG-1 all from the heterologous *mec-3* promoter and found that the expression of TRN markers was turned off, suggesting that EGL-44/EGL-46 can repress TRN fate in the presence of ZAG-1 (Figure 4F). These data suggest that the mutual inhibition between ZAG-1 and EGL-44/EGL-46 occurs only at the transcription level and not likely through direct interaction. Consistent with this idea, ZAG-1 failed to interact with either EGL-44 or EGL-46 in yeast two-hybrid assays (MEC-3 also failed to interact with EGL-44, EGL-46, and ZAG-1; Figure S3).

We also tested the touch sensitivity of the above transgenic animals and the functional results are consistent with the marker expression results. Animals carrying the *uIs211[mec-3p::egl-44]* transgene were completely insensitive to gentle touch; co-expression of *zag-1* from *mec-3* promoter restored the sensitivity in animals where the TRN markers were reactivated but failed to do so when *mec-3p::egl-46* was also co-expressed (Figure S5).

### EGL-44/EGL-46 and ZAG-1 regulate a switch between FLP and TRN fates

We next investigated how EGL-44/EGL-46 and ZAG-1 regulated FLP fate using several genes (the stomatin gene *sto-5*, the glycoprotein gene *dma-1*, the dynein regulator *bicd-1*, and the FMRFamide-like peptide gene *flp-4*) that were expressed in FLP neurons (Aguirre-Chen et al., 2011; Kim and Li, 2004; Liu and Shen, 2011) but not the TRNs (Figure 5A). FLP expression of all four genes depended on EGL-44 and EGL-46 (Figure 5B and C). Moreover, the expression of the FLP and TRN markers was mutually exclusive; FLP neurons in *egl-44* and *egl-46* mutants or animals carrying *mec-3p::zag-1* transgene never showed mixed expression of the FLP- and TRN-specific genes (Figure 5E). Mutation in *zag-1* or the misexpression of *egl-44* in TRNs activates the FLP markers in addition to turning off the TRN markers (Figure 5B and S6), and we never observed mixed expression of both types of markers (Figure 5E). Since FLP markers were not expressed in *egl-44; zag-1* double mutants, our results suggest that EGL-44/EGL-46 not only repressed the TRN fate but also promoted the FLP fate. Thus, the action of EGL-44/EGL-46 and ZAG-1 results in two mutually exclusive fates.

**Figure 5.**
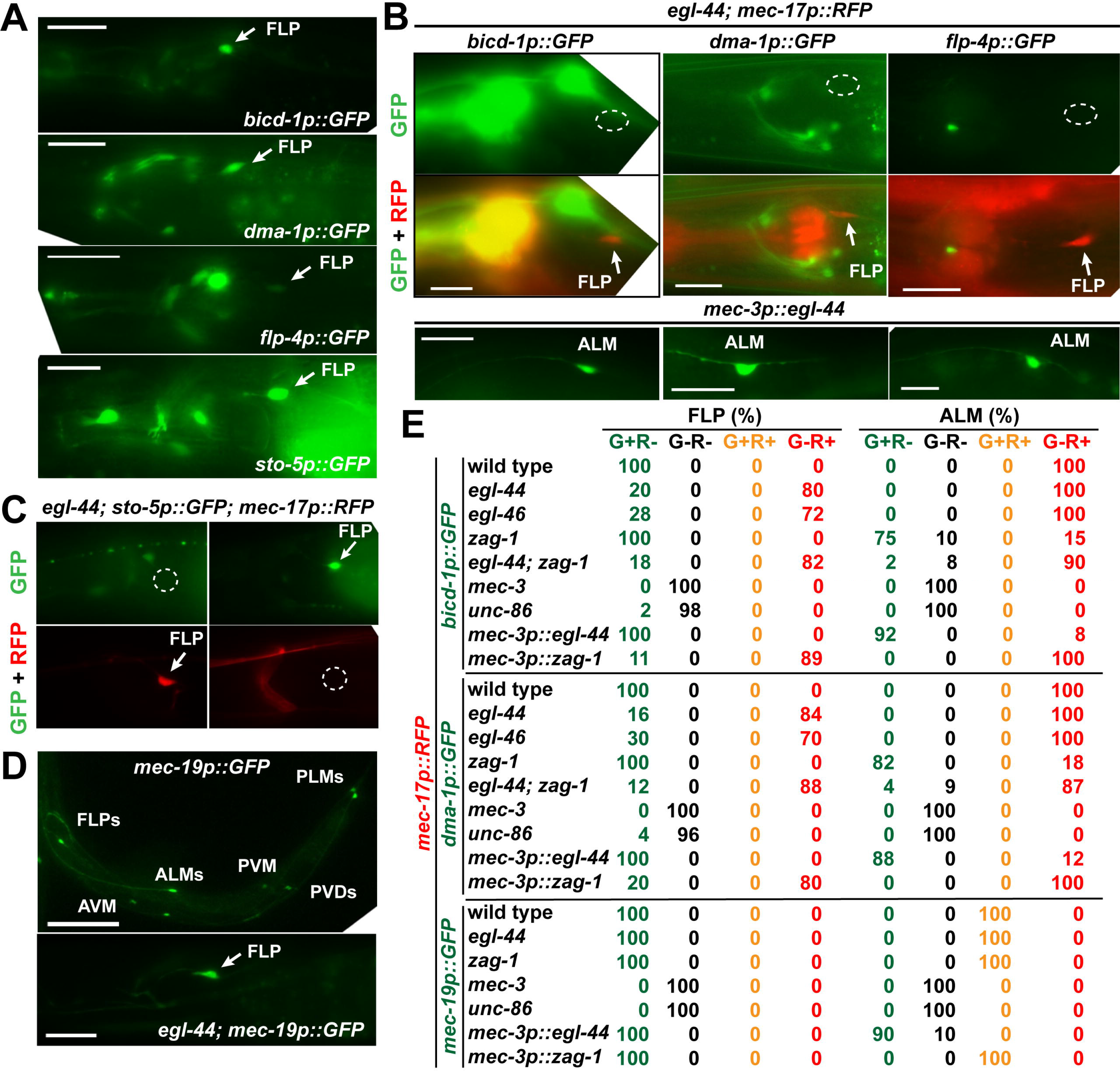
EGL-44/EGL-46 simultaneously induces FLP genes and suppresses TRN genes. (A-B) Various FLP fate reporters were expressed in wild-type FLP neurons but not in *egl-44* mutants. (C) The expression of FLP and TRN fate markers was mutually exclusive in FLP neurons of *egl-44* mutants. (D) The expression of *mec-19* was not affected by mutations in *egl-44*. (E) Percentage of FLP and ALM neurons expressing the green (G) and red (R) markers in various strains. The number of cells > 40.

The EGL-44/EGL-46 ZAG-1 bistable switch affected neuronal morphology as well as transcriptional reporters. For example, *egl-44* misexpression in ALM neurons resulted in an ectopic posterior neurite characteristic of FLP neurons (Figure S7), and *zag-1* expression in FLP and PVD neurons drastically reduced the number of dendritic branches (Figure S7). This latter result is consistent with ZAG-1’s role in preventing PVM from adopting a PVD-like morphology (Smith et al., 2013).

*dma-1*, *bicd-1*, and *flp-4* were also expressed in PVD neurons, suggesting that FLP and PVD share a common genetic program. The expression of these PVD markers was blocked by the misexpression of ZAG-1 in PVD neurons, suggesting that ZAG-1 inhibited the unknown PVD fate regulator (repressor). On the other hand, the *sto-5* reporter was expressed in FLP but not PVD neurons, indicating differences in the transcriptome of the two cell types (Figure S8).

The activation of the FLP and PVD markers, like the TRN markers, required UNC-86 and MEC-3, which are the common terminal selectors for the three fates (Figure 5E and S6B). We identified one common terminal differentiation gene, *mec-19*, which was expressed in TRN, FLP, and PVD neurons at similar levels (and no other cells); its expression depended on UNC-86 and MEC-3 but was not affected by the loss of EGL-44/EGL-46 and ZAG-1 (Figure 5D-E). Thus, *unc-86* and *mec-3* serve as the “ground-state selectors” that promote a common ground state shared by the three types of neurons, and *mec-19* is a marker for this ground state. Subsequently, *zag-1* and *egl-44* act as “modulators” that shift the ground state towards specific fates. Therefore, the development of a particular fate requires the activities of both the ground-state selectors and the fate-restricting modulators. Our results are consistent with the findings in *C. elegans* cholinergic motor neurons, where ground-state markers, such as *cho-1* (choline transporter), *cha-1* (choline acetyltransferase), and *unc-17* (vesicle acetylcholine transporter), are activated by selectors and not regulated by subtype-specific repressors (Kerk et al., 2017).

Among the six TRNs, ALM maintains the default TRN state, whereas PLM undergoes further differentiation controlled by the posterior Hox protein EGL-5, which represses UNC-86/MEC-3-dependent ALM genes and activates UNC-86/MEC-3-independent PLM genes (Zheng et al., 2015a). We found that the ALM gene *mir-84*, but not the PLM gene *rfip-1*, was expressed in the FLP neurons of *egl-44* mutants (FLP neurons do not express EGL-5; Zheng et al., 2015b; Figure S9). These results suggest that that the bistable switch controls the general TRN fate not TRN subtype identity.

### EGL-44/EGL-46 inhibits the expression of TRN genes by binding to UNC-86/MEC-3-binding sites and by suppressing ALR-1 in FLP neurons

We next investigated the mechanism, by which EGL-44/EGL-46 prevented the expression of TRN terminal differentiation genes. Although we found two UNC-86/MEC-3 binding sites required for the expression of a minimal TRN-specific promoter (a 184-bp *mec-18* promoter) in TRNs, mutational analysis of this minimal promoter failed to identify any additional, discrete *cis*-regulatory element that mediated its repression in FLP neurons (Figure S10). Thus, EGL-44/EGL-46 seems unlikely to suppress TRN markers *via* repressive elements that are distinct from the UNC-86/MEC-3 binding sites. We then tested the possibility that EGL-44/EGL-46 acts through the UNC-86/MEC-3 binding site, since EGL-44 belongs to the TEA domain class transcription factors, which recognize DNA sequences similar to the UNC-86/MEC-3 binding site (Figure 6A) (Jiang et al., 2000; Zhang et al., 2002). EGL-44 bound to previously identified UNC-86/MEC-3 motifs in the *mec-4*, *mec-7*, *mec-17*, and *mec-18* promoters in electrophoretic mobility shift assays (Figure 6B), suggesting that EGL-44 directly contacts the TRN promoters. The EGL-44/EGL-46 complex also bound to the same EGL-44-binding motif in the *mec-4* promoter, suggesting EGL-46 may act as a corepressor (Figure 6C). The association of EGL-44/EGL-46 to the *cis*-regulatory element bound by UNC-86/MEC-3 suggests that EGL-44/EGL-46 in FLP neurons may prevent the activation of TRN genes by occluding the UNC-86/MEC-3-binding sites essential for the expression of TRN fate.

**Figure 6.**
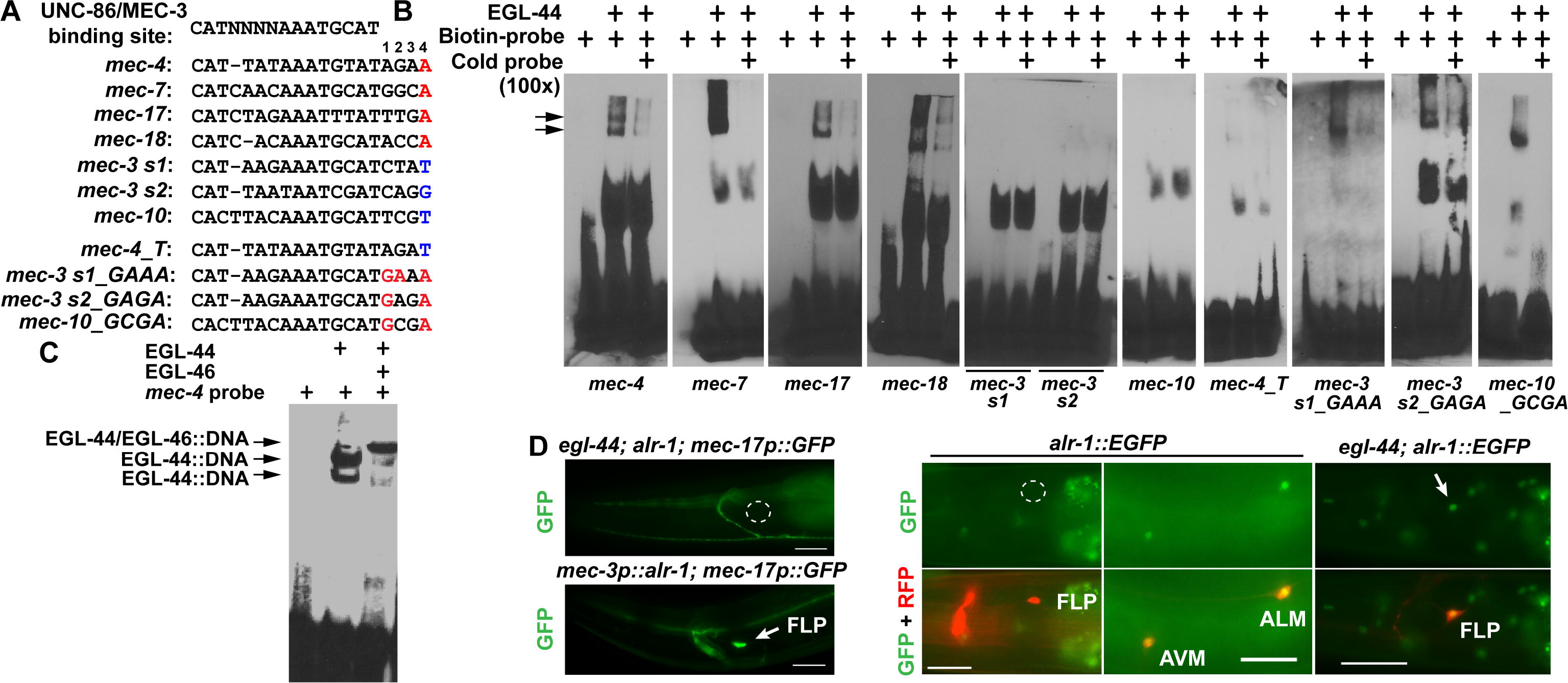
EGL-44/EGL-46 inhibits the expression of TRN genes by binding to the same *cis-*regulatory elements bound by UNC-86/MEC-3. (A) The alignment of various *cis*-regulatory motifs tested in electrophoretic mobility shift (EMSA) assays. Positions 1-4 were assigned to the four nucleotides following the consensus UNC-86/MEC-3 binding site (Zhang et al., 2002). Nucleotides in red were considered important for EGL-44 binding and nucleotides in blue were responsible for the lack of EGL-44 binding. (B) The binding of recombinant EGL-44 proteins to various probes in EMSA assays. Arrows point to the band of EGL-44::DNA complexes. (C) The binding of EGL-44/EGL-46 to the *mec-4* probes. (D) The expression of TRN fate markers in *egl-44; alr-1* double mutants and animals misexpressing ALR-1 from the *mec-3* promoter and the expression of a fosmid-based *alr-1* reporter *wgIs200[alr-1::EGFP]* in wild-type animals and *egl-44* mutants.

**Figure 7.**
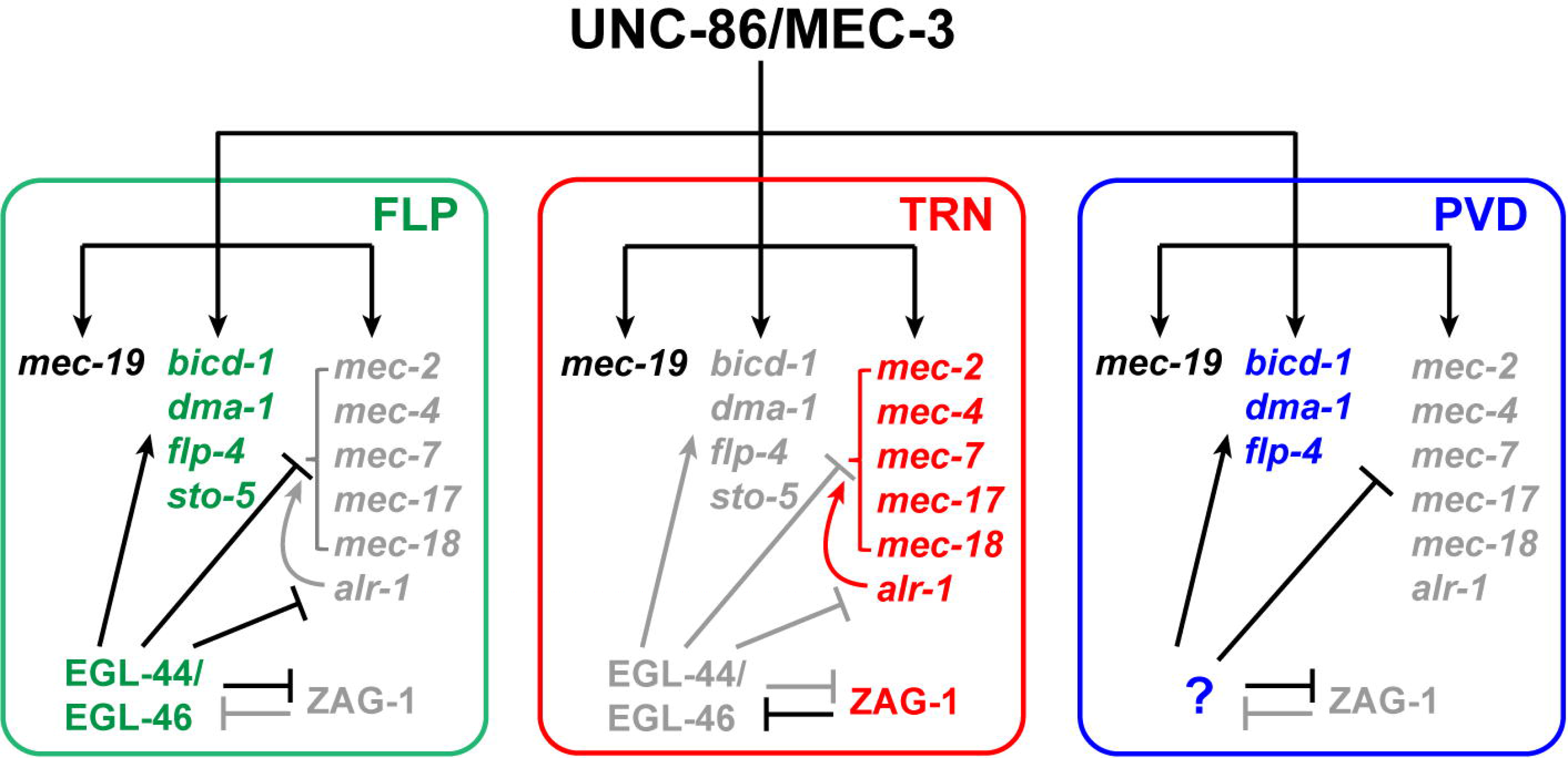
A model for the regulatory mechanisms controlling the cell fate specification among FLP, PVD, and TRN fates. Gene names in green, red, and blue indicate genes expressed in FLP, TRN, and PVD neurons, respectively; genes whose names are black were commonly expressed in all three types and those whose names are grey were repressed.

How EGL-44/EGL-46, however, avoids repressing UNC-86/MEC-3 targets that are commonly expressed in FLP and TRNs is unclear. One such target is *mec-3* itself, whose activation and maintenance depend on two UNC-86/MEC-3 binding sites (Xue et al., 1992). We found that the two sites were not bound by EGL-44 (Figure 6B), which explains why *mec-3* expression is not affected by the presence of EGL-44/EGL-46 in FLP neurons. Comparing the *cis*-regulatory motif sequences on the *mec-3* promoter with those on TRN-specific promoters, we found that the four nucleotides (positions 1 to 4 in Figure 6A) following the UNC-86/MEC-3 binding sequence showed significant divergence between EGL-44-binding and nonbinding sites. In particular, the EGL-44-binding sites all contain adenine at the fourth position and the nonbinding sites do not (Figure 6A). Changing this adenine to thymine eliminated the binding of EGL-44 to the site on the *mec-4* promoter (Figure 6B). However, converting other nucleotides to adenine at the fourth position was not sufficient to enable the binding of EGL-44 to *mec-3* promoter motifs and coordinated change of the nucleotides on the first and second positions are needed to evoke EGL-44 binding (Figure 6B). Similar results were also obtained for the UNC-86/MEC-3 target gene *mec-10*, which is expressed in both TRN and FLP cells (Huang and Chalfie, 1994). The UNC-86/MEC-3 site on *mec-10* promoter was not bound by EGL-44 and changing the nucleotides on the first and fourth position enabled EGL-44 binding. Thus, our results suggest that EGL-44/EGL-46 can differentiate TRN/FLP common genes from TRN-specific genes *via* the *cis*-regulatory sequences in their promoters.

We next tested whether converting the EGL-44 binding site to a nonbinding site in a TRN promoter could prevent the suppression by EGL-44/EGL-46 in FLP neurons *in vivo*. Contrary to our expectation, a *mec-4* promoter reporter harboring the adenine-to-thymine change at the fourth position was not activated in FLP neurons, although its TRN expression was preserved. In the 184-bp *mec-18* promoter, changing the four nucleotides following the UNC-86/MEC-3 binding sequence from ACCA to CTAT completely abolished EGL-44 binding *in vitro*; but the mutant reporter was only weakly expressed in ~30% of FLP neurons and remained silenced in the rest ~70% (Figure S11). In addition, creating an EGL-44-binding site in the *mec-3* promoter by mutating the regulatory motif did not suppress *mec-3* expression in FLP neurons (Figure S11A). The discrepancy between the *in vitro* and *in vivo* results suggests that the lack of EGL-44 binding to TRN differentiation genes *per se* is not sufficient to distinguish TRN fate from the TRN/FLP ground state. In addition to directly binding to the *cis*-regulatory elements of TRN promoters, EGL-44/EGL-46 may also inhibit TRN genes by activating or repressing the expression of other *trans*-acting factors.

One such factor may be ALR-1, which is an ortholog of human Arx and Drosophila *aristaless* and is needed for the robust differentiation of TRN fate (Topalidou et al., 2011). Loss of *alr-1* variably reduced but did not eliminate the expression of TRN markers in the TRNs in wild type and strongly eliminated the ectopic expression of TRN markers in FLP neurons in *egl-44* and *egl-46* mutants (Figure 6D and S12A). This difference between reduction and elimination may result from the fact that FLP neurons have lower *mec-3* expression than the TRNs (Topalidou et al., 2011) and thus a stronger need for ALR-1 to activate TRN markers. The observation that *alr-1* is epistatic to *egl-44* and *egl-46* suggested that ALR-1 is a downstream effector suppressed by EGL-44/EGL-46. Indeed, a fosmid-based *alr-1* translational reporter, which was normally expressed in TRNs but not FLP neurons, became de-repressed in FLP cells in *egl-44* and *egl-46* mutants (Figure 6D and S12A).

The lack of ALR-1 and the lower level of *mec-3* in wild-type FLP neurons may explain the inactivation of the mutated TRN fate reporters that cannot be bound by EGL-44/EGL-46. Supporting this hypothesis, forced expression of ALR-1 in FLP neurons ectopically activated even the wild-type TRN fate reporters (Topalidou et al., 2011) (Figure 6D), suggesting that ALR-1 not only promotes TRN fate but also overcomes the direct suppression from EGL-44/EGL-46 on the TRN genes. This ability of ALR-1 may result from ALR-1 upregulating *mec-3* (Topalidou et al., 2011) and directly interacting with TRN promoters (ChIP-seq data; Figure S12B). Therefore, EGL-44/EGL-46 inhibits TRN-specific genes by both occupying the UNC-86/MEC-3 site in the TRN promoters and suppressing TRN fate-promoting transcription factor ALR-1. Consistent with this model, overexpression of MEC-3 in FLP neurons could overcome EGL-44/EGL-46-mediated inhibition and activate the TRN program (Topalidou and Chalfie, 2011) presumably by both retaking the UNC-86/MEC-3 sites and by activating *alr-1*, which is a *mec-3*-dependent gene (Topalidou et al., 2011).

### Spatial and temporal expression of *zag-1* and *egl-44* are mutually exclusive

Given the mutual inhibition of *zag-1* and *egl-44* in the FLP neurons and the TRNs, we asked whether the two genes were generally expressed in different cells in the nervous system. Using the neurotransmitter maps for glutamatergic, cholinergic, and GABAergic neurons (Gendrel et al., 2016; Pereira et al., 2015; Serrano-Saiz et al., 2013), we found that, in addition to TRNs, *zag-1* was expressed in the AIB, AIM, AIN, AIZ, AVA, AVB, AVD, AVE, AVG, AVK, AVL, M4, M5, RIA, RIB, RIF, RIG, RIM, RIV, RMD, RME, RMF, RMH, SIA, and SMD neurons in the head, all the DD, VD, and VC neurons in the ventral cord, and the DVA, DVB, LUA, PDA, PVC, PVP, PVQ, PVR, and PVT neurons in the tail. *zag-1* is also expressed in the serotonergic HSN neurons. In comparison, *egl-44* expression was much more restricted, being found only in the FLP, ADL, and SAB neurons, as well as a few VA and VB motor neurons. Moreover, *egl-44* was widely expressed in hypodermis, pharynx, and intestine, whereas *zag-1* expression was absent in these tissues. We also constructed an *egl-44::RFP* reporter and crossed it with *zag-1:EGFP* and did not observe overlapping expression in any cell (Figure S13A). The above data support the hypothesis that *zag-1* and *egl-44* expression are mutually exclusive in the nervous system and throughout all tissues of the animal. Given the mutual inhibition of EGL-44 and ZAG-1 in the TRNs/FLPs, we expected that the loss of *zag-1* might affect *egl-*44 expression more broadly. We did not, however, observe a systematic upregulation of *egl-44::GFP* expression in the nervous system in either the *zag-1(zd86)* hypomorphic mutants or the arrested L1 animals of *zag-1(hd16)* null mutants (Figure S13B and C). Similarly, tissues like hypodermis, pharynx and intestine did not gain *zag-1* expression upon the loss of *egl-44* either, suggesting that, in addition to the reciprocal inhibition, other activating signals are needed to create the expression pattern of the two transcription factors.

A few neurons expressed both *zag-1* and *egl-44* but did so at different times. *egl-44* and *egl-46* were expressed in the precursors of the postembryonic TRNs (the AVM and PVM neurons) and persisted in these cells for a few hours after their generation (Feng et al., 2013; Wu et al., 2001), but they were not expressed in terminally differentiated AVM and PVM. These differentiated cells expressed *zag-1*, which promoted the TRN fate. The transient expression of *egl-44* and *egl-46* may be to temporarily block selector functions and to ensure the correct timing of differentiation. Similarly, *egl-44* and *egl-46* were transiently expressed in the early embryos in HSN neurons before they migrated and differentiated (Wu et al., 2001) (Figure S13D), whereas *zag-1* was expressed in the terminally differentiated HSN neurons in adults and was required for the activation of the HSN fate marker *tph-1p::GFP* (Figure S13E and F).

## Discussion

### Using RNAi to identify genes involved in cell fate determination

We demonstrate here that a systematic RNAi screen using a library of transcription factors can identify neuronal cell fate regulators, particularly those genes whose mutations lead to lethality or sterility. Our previous forward genetic screens, which searched for viable mutants with touch-sensing defects, despite reaching saturation, only identified *unc-86* and *mec-3* as the TRN fate determinants (Chalfie and Au, 1989; Chalfie and Sulston, 1981). Using the RNAi screen, we not only recovered *unc-86* and *mec-3* blindly, but also identified *ceh-20*, *ldb-1*, and *zag-1* as genes required for the expression of TRN fate. These latter genes would not have been identified in our previous screens, because null mutations in them lead to early larval arrest. In particular, our RNAi screen yielded ZAG-1 as a new TRN fate determinant and the third piece in a combinatorial code (UNC-86, MEC-3, and ZAG-1) that defines TRN fate.

Our experiments raise two concerns about the reliability of such RNAi screens. First, only 5 of the 14 genes identified from the screen were confirmed with mutants, and the remaining 9 genes (65%) appeared to be false positives. A similar problem was encountered in a genome-wide RNAi screen for genes involved in the specification of ASE neuron in *C. elegans* (Poole et al., 2011). More recently, Kok et al. (2015) found that approximately 80% of morpholino-induced phenotypes in zebrafish embryos were not observed in mutants. Similar discrepancies between RNAi and mutant phenotypes were also observed in mice (Daude et al., 2012) and Arabidopsis (Gao et al., 2015). Such widespread differences may be attributed to the mistargeting of dsRNAs or to genetic compensation (the upregulation of a network of genes that compensate for genetic loss but not knockdown; Rossi et al., 2015). Second, our screen failed to recover *ceh-13*, which is known to affect the TRN fate in ALM neurons (Zheng et al., 2015b), suggesting the existence of false negatives. In addition, because the PLM cell body is located in the tail region (posterior to the intestine), the bias of feeding RNAi towards more anterior cells led to the failure of recovering *egl-5*, whose loss only affects PLM differentiation (Zheng et al., 2015b).

### ZAG-1 prevents the inappropriate expression of repressors and safeguards cell fate determination

In this study, we found that neurons develop protective mechanisms to prevent the improper activation of repressors and to ensure the differentiation of the correct cell fate. Specifically, ZAG-1 prevents the expression of repressors EGL-44 and EGL-46 in the TRNs. Because the EGL-44/EGL-46 complex is a powerful inhibitor of TRN fate (its derepression in *zag-1* mutants or misexpression in TRNs can completely shut off the expression of TRN genes), the function of ZAG-1 is essential to ensuring the specification of TRN fate.

As a differentiation regulator, ZAG-1 is required for the adoption of TRN fate but does not directly bind to the *cis*-regulatory elements within TRN promoters and so does not act as a selector. Moreover, unlike the highly confined expression of terminal selectors and repressors (e.g. both *mec-3* and *egl-44* are expressed in few neurons), *zag-1* is expressed in many neurons in the head and tail ganglia and ventral cord motor neurons, as well as various muscles (this study; Wacker et al., 2003). Thus, ZAG-1, as a transcriptional repressor, may serve as a cell fate protector for many different neurons. Already ZAG-1 is found to be required for the fate specification of at least TRN, HSN, PVQ, and M4 neurons (this study and Clark and Chiu, 2003; Ramakrishnan and Okkema, 2014), although whether ZAG-1 also regulates HSN, PVQ, and M4 fates by preventing repressor expression is unclear.

The widespread expression of *zag-1* in neurons is also consistent with its function in controlling neurite outgrowth (this study; Clark and Chiu, 2003; Wacker et al., 2003). This separate function, observed in many different neurons, appears to be independent of its role in inhibiting the expression of repressors in cell-fate specification.

### Diverse mechanisms of repressor function

Repressors affect effector gene expression in several ways. Repressors can act through discrete *cis*-regulatory elements that are close to but separate from the selector binding site on the promoter of effector genes (Kerk et al., 2017), and the mechanism of repression may involve histone modification and the recruitment of histone deacetylase (Winnier et al., 1999). Our study suggests that repressors (EGL-44/EGL-46) may directly occupy the selector binding site (for UNC-86/MEC-3) and, thus, could prevent selector binding.

Repressors that inhibit effector genes directly can also activate effector genes indirectly by inhibiting other repressors (Kerk et al., 2017). EGL-44/EGL-46 does activate FLP genes in addition to inhibiting TRN genes, whether it does so by repressing other repressors is unclear.

All of these methods shape the final collection of effector genes that define a particular cell fate. ZAG-1, however, appears to act differently because it does not interact with effector genes directly (we did not find any UNC-86/MEC-3-induced effector gene that was directly repressed by ZAG-1). Instead, a major function for ZAG-1 in TRNs is to prevent the expression of repressors, including EGL-44/EGL-46 and possibly the unknown repressor that inhibits TRN fate in PVD neurons. Thus, ZAG-1 may be devoted to inhibiting the alternative FLP and PVD fates, while UNC-86/MEC-3 induces TRN fate. The uncoupling between the two modules suggests that they may have evolved independently. We distinguish the action of EGL-44/EGL-46, which prevents downstream gene expression, from that of ZAG-1, which regulates *egl-44* and *egl-46* expression, by calling the former a repressor and the latter an inhibitor.

### Negative feedback loop controls binary fate choice and neuronal diversification

The repressors reported by Kerk et al. (2017) formed a hierarchy that regulated effector gene expression in motor neurons. In this study, however, we showed that by downregulating each other’s gene expression, a repressor and an inhibitor can form a bistable switch that ensures proper cell differentiation. The diversification of three types of sensory neurons (TRNs, FLP, and PVD neurons) that share the same selectors (UNC-86 and MEC-3) use a set of binary fate choices. The TRN versus FLP fate choice is controlled by the negative feedback loop between ZAG-1 and EGL-44, while the TRN versus PVD choice might be similarly controlled by mutual inhibition between ZAG-1 and an unknown repressor that helps specify PVD fate and represses TRN fate.

In the absence of both components of the switch (*egl-44; zag-1* mutants), both TRNs and FLP neurons expressed the TRN genetic program, suggesting that the ground state is a TRN-like fate and the selectors UNC-86/MEC-3 activate TRN genes by default. EGL-44/EGL-46-induced modification of the default genetic program gives rise to the FLP fate. Our results are different from the observation in motor neurons, where the loss of repressors leads to a “mixed ground state” that is not similar to any of the five motor neuron types (Kerk et al., 2017). This distinction may result because ZAG-1 only inhibits EGL-44 and does not directly regulate effector genes, whereas in motor neurons all repressors interact with effector genes.

The choosing of one of two alternative fates appears to be a common way to diversify neuronal types and subtypes. Notable examples include the fate choices between Drosophila R7 and R8 photoreceptors (Mikeladze-Dvali et al., 2005) and between the left and right ASE gustatory neurons in *C. elegans* (Sarin et al., 2007); both decisions are mediated by bistable feedback loops. In vertebrates, mutual repression between Hox proteins directs motor neuron diversification in mouse spinal cord (Dasen et al., 2003; Dasen et al., 2005), and the neurotransmitter identity of cortical neurons is also subjected to binary regulation (Lodato et al., 2014; Nakatani et al., 2007).

The function of ZEB family transcription factors in regulating such binary cell fate choices appears to be evolutionary conserved. The Drosophila homolog of ZAG-1, Zfh1, promotes a GW motor neuron fate over an EW interneuron fate in the 7-3 neuroblast lineage, and its expression is suppressed by the steroid receptor family transcription factor Eagle in the EW interneuron (Lee and Lundell, 2007); whether Zfh1 also suppressed Eagle in GW motor neuron is unclear. In another example, the mutual antagonism between Zfh1 and another zinc-finger transcription factor, Lame duck, regulates the decision between pericardial cell and fusion competent myoblast fates in the mesoderm (Sellin et al., 2009). Although a genetic interaction between Zfh1 and the Drosophila EGL-44, Scalloped, has not been reported, these results suggest that Zfh1 can form regulatory switches with other transcription factors. Mouse homologs of ZAG-1, ZEB1 and ZEB2, repress tissue differentiation during early embryogenesis, induce epithelial mesenchymal transition (EMT), and are essential for neural tube development (Vandewalle et al., 2009). Their roles in specifying terminal cell fates, however, are unclear, because their knockout leads to embryonic lethality (Miyoshi et al., 2006).

### Binary fate switches may enable the generation of neuronal diversity

Because the introduction of self-reinforcing, bistable switches can generate diversity within pre-existing neuronal fates, we envision that the extraordinary variety of neuronal types in the nervous system evolved from a few primitive neuronal fates through stepwise addition of binary switches. For example, since TRN, FLP, and PVD fates all require the same selectors, UNC-86 and MEC-3, they may be derived from a common ancestral fate. In fact, the existence of UNC-86/MEC-3 target genes like *mec-19*, which is expressed in all the FLP, PVD, and TRNs but not any other neuron, suggests such an ancestral fate.

One possible evolutionary derivation of these cells is that some ancestral TRN-like cells acquired the ability to express *egl-44* and *egl-46*. This expression led to the suppression of TRN genes, the activation of FLP genes, and the emergence of the FLP neuron type. The fact that the ground state is a TRN-like state seems to support this hypothesis. Moreover, since *egl-44* is primarily expressed in non-neuronal tissues, mutations in the regulatory elements of *egl-44* might have allowed expression in some neurons. Thus, EGL-44 may have been co-opted to induce divergence among neurons that share a common fate, and this cell fate divergence was subsequently stabilized by the establishment of a negative feedback loop between EGL-44 and ZAG-1. Alternatively, a regulatory element in the *zag-1* gene, which was already present in the ancestral TRNs to repress *egl-44* and *egl-46*, may have been mutated to prevent its expression in some ancestral TRN cells. This loss led to the de-repression of *egl-44* and *egl-46* and the subsequent acquisition of the FLP fate.

Overall, we imagine that a limited number of ground-state selectors may first define a handful of shared states, and subsequently a series of binary fate switches carry out further differentiation that modifies the ground state to generate a diverse array of terminal neuronal fates. The broad expression of ZAG-1 in the nervous system and the lack of increased EGL-44 expression in *zag-1* mutants suggests that ZAG-1 may form bistable regulatory loops with many different inhibitors. Moreover, the fact that ZAG-1 expression is only found in a subset of neurons that express the same selector supports the hypothesis that ZAG-1 may ensure diversification from a shared ground state by inhibiting repressors; e.g. *zag-1* is expressed in 7/17, 6/15, and 3/11 classes of neurons that use UNC-86, UNC-3, and CEH-14 as a terminal selector, respectively (classification according to Hobert, 2016). Selective loss of *zag-1* expression in some cells may prevent these cells from committing to particular fates and allowing them to adopt alternative ones. Finally, the broad expression of ZEB family transcription factors in the nervous system of both Drosophila (Lai et al., 1991) and mice (Vandewalle et al., 2009) suggests possibly a conserved role for them in regulating binary fate choices and the generation of neuronal diversity.

## Materials and Methods

### Strains, Constructs, and transgenes

*C. elegans* wild type (N2) and mutant strains were maintained as previously described (Brenner, 1974). Most strains were provided by the *Caenorhabditis* Genetics Center, which is funded by NIH Office of Research Infrastructure Programs (P40 OD010440), or the National BioResource Project of Japan. VH514, *zag-1(hd16)/unc-17(e113) dpy-13(e184)* was used as the balanced null allele for *zag-1* and *zag-1(zd86)* was used as a hypomorphic allele. For other genes identified from the RNAi screen (see below), we tested *zip-4(tm1359)*, *nhr-119(gk136908)*, *nhr-166(gk613)*, *nhr-159(tm2323)*, *egl-38(ok3510)/nT1[qIs51]*, *hmbx-1(ok3467)*, *fkh-2(ok683)*, *lin-40(ku285)*, *lin-40(s1506) unc-46(e177)/eT1*, *elt-6(gk723)*, and *elt-6(gk754)*. Other mutant alleles used in this study include *ahr-1(ju145)*, *egl-44(n1080)*, *egl-46(n1127)*, *mec-3(u184)*, and *unc-86(u5)*.

A 2.4 kb *mec-3* promoter, a 2.2 kb *zag-1* promoter, a 2.2 kb *bicd-1* promoter, 4.9 kb *dma-1* promoter, 3.1 kb *flp-4* promoter, 2.3 kb *sto-5* promoter, and 1.3 kb *mec-19* promoter were cloned from wild type (N2) genomic DNA into the Gateway pDONR221 P4-P1r vector. The genomic coding region of GFP, *zag-1*, *egl-44*, *egl-46*, *alr-1*, and *ahr-1* were cloned into Gateway pDONR221. The resulted entry vectors, together with pENTR-*unc-54*-3’ UTR and the destination vector pDEST-R4-R3 were used in the LR reaction to create the final expression vectors. Gateway cloning was performed according to the manual provided by Life Technologies (Grand Island, NY). To generate transgenic animals, we injected DNA constructs (5 ng/l for each expression vector) into the animals to establish stable lines carrying an extrachromosomal array; at least three independent lines were tested. In some cases, the transgene was integrated into the genome using -irradiation (Mello et al., 1991), and at least three integrant lines were outcrossed and examined. For the *mec-3p::zag-1; mec-3p::egl-44; mec-3p::egl-46* triple transgenic animals, the injection was repeated twice, and three stable lines were obtained from each injection. Similar results were obtained from the replicates.

DNA constructs TU#625 and TU#626 (Wu et al., 2001) contain translational GFP fusion of *egl-44* and *egl-46*, respectively, and were injected into animals to form reporters for the two genes, named as *uIs215[egl-44::GFP]* and *uEx927[egl-46::GFP]*. TU#924 contains a 400 bp *mec-18* promoter inserted into pPD95.75 between HindIII and BamHI sites, and this *mec-18p::GFP* construct was used as a template to create a series of promoter variants shown in Figure S10 using the Q5 site-directed mutagenesis kit from New England Biolabs (Ipswich, MA).

Transgenes *zdIs5[mec-4p::GFP]* I, *muIs32[mec-7p::GFP]* II, *uIs31[mec-17p::GFP]* III, *uIs115[mec-17p::RFP]* IV, *uIs134[mec-17p::RFP]* V, and *uIs72[mec-18p::mec-18::GFP]* were used as fluorescent markers for the TRN cell fate. *uEx1104[Pbicd-1::GFP]*, *uEx1105[dma-1p::GFP]*, *uEx1106[flp-4p::GFP]*, and *uIs232[sto-5p::GFP]* served as FLP fate markers. *zdIs13[tph-1p::GFP]* and *vsIs97[tph-1p::DsRed2]* were used as HSN fate marker. *uIs22[mec-3p::GFP]* and *uIs152[mec-3p::RFP]* were used as *mec-3* transcriptional reporter; *uEx1007[mec-3p::mec-3::GFP]* and *wgIs55[mec-3::TY1::EGFP::3xFLAG]* were used as *mec-3* translational reporter. *wgIs83[zag-1::TY1::EGFP::3xFLAG]*, *wgIs476[unc-86::TY1::EGFP::3xFLAG]*, *wgIs200[alr-1::TY1::EGFP::3xFLAG]*, *leEx1709[ahr-1::GFP]* and *uEx1107[mec-19p::GFP]* served as the reporters for *zag-1*, *unc-86*, *alr-1*, *ahr-1*, and *mec-19* respectively. *uIs211[mec-3p::egl-44]*, *uEx926[mec-3p::zag-1]*, and *uEx1027[mec-3p::alr-1]* were used for misexpression.

To perform cell identification, we crossed *wgIs83[zag-1::TY1::EGFP::3xFLAG]* and *uIs215[egl-44::GFP]* into the *otIs518[eat-4::SL2::mCherry::H2B]*, *otIs544[cho-1::SL2::mCherry::H2B]*, and *otIs564 [unc-47::SL2::H2B::mChopti]*, labeling glutamatergic, cholinergic, and GABAergic neurons (Gendrel et al., 2016; Pereira et al., 2015; Serrano-Saiz et al., 2013), respectively. We identified *zag-1* and *egl-44* expressing neurons, based on the position and neurotransmitter identity of the cells expressing GFP.

### RNAi screen

RNAi screen was performed using a modified bacteria-feeding protocol previously reported (Kamath et al., 2003; Poole et al., 2011). We used the Ahringer RNAi library from Source Bioscience (http://www.lifesciences.sourcebioscience.com/) and the list of 392 RNAi clones targeting transcription factors were generated by searching WormBase WS238 using Gene Ontology terms related to “DNA binding” and “transcription factor activity” (see the complete list in Table S1). To perform the RNAi experiments, we seeded bacteria expressing dsRNA on NGM agar plates containing 6 mM IPTG and 100 µg/ml ampicillin. One day later, eggs from TU4429, *eri-1 (mg366); lin-15B (n744); uIs134[mec-17p::RFP]* animals were placed onto these plates; the eggs hatched and grew to adults at 20°C. The F1 progeny of these worms were scored for the expression of TRN markers at the second larval stage. RNAi clones were considered to be positive if more than 15% (n > 20) of the treated animals failed to show RFP expression in the ALM neurons in at least two of the three replicate plates. Three initial rounds of screens were performed on all the 392 clones, and 14 clones were found positive in all the three rounds. Two more screening rounds were then conducted on these 14 RNAi clones, which were all confirmed to be positive. We sequenced the inserts of all positive clones to confirm the identity of the target genes.

### Yeast two-hybrid assay

Yeast media and plates were prepared according to recipes from Clontech (Mountain View, CA) and yeasts were grown at 30°C. The yeast strain PJ69-4a (provided by Songtao Jia at Columbia University) used for the two-hybrid assays contains GAL1-HIS3, GAL2-ADE2, and GAL7-lacZ reporters. Vectors pGAD424 and pGBT9 (Clontech) were used to express proteins fused to the yeast activating domain (AD) and binding domain (BD), respectively. cDNA fragments of *mec-3*, *zag-1*, *ldb-1*, *egl-44*, and *egl-46* were cloned into the two-hybrid vectors either using restriction enzymes or with Gibson Assembly (NEB).

Combinations of the AD or BD vectors were co-transformed into yeast using the Frozen-EZ II kit from Zymo Research (Irvine, CA) and using empty vectors as negative controls. Growth assays were performed by growing individual colonies overnight in selective media lacking tryptophan and leucine. Cultures were then diluted to let OD600 become 0.5, and 10 l of a further 1:10 diluted culture were spotted onto plates lacking histidine to test the expression of the HIS3 reporter. Plates were imaged after 2 days of growth. Liquid β-galactosidase assays were performed using the Yeast β-Galactosidase Assay Kit (Thermo Scientific, Rockford, IL).

### Electrophoretic mobility shift assay (EMSA)

Recombinant GST::EGL-44 proteins were produced in *E. coli* BL21 (DE3) using the expression vector pGEX-6p-1 (Amersham Pharmacia Biotech, UK) and purified using affinity chromatography columns filled with Glutathione Sepharose 4B beads (Amersham). EGL-44 was cleaved off the column using PreScission Protease (Amersham). EGL-46 was expressed using the pET32a vector (Novagen, Madison, WI) and purified using the S-Tag rEK purification kit (Novagen).

Gel mobility shift assays were performed using a modified protocol previously reported (Xue et al., 1993). DNA probes were labeled with Biotin by Biotin 3’ End labeling kit (Pierce, Rockford, IL) and then annealed into double strands. 100 ng of proteins were incubated with 20 fmol (0.5~0.7 ng) probe at room temperature for 30 min, and the mixture was loaded onto a 10% TBE polyacrylamide mini gel (Bio-Rad, Hercules, CA). 2 pmol unlabeled probe (100x) was added for the cold probe competition. The gel was transferred to a 0.2 mm nylon membrane, which was then treated with UV light to crosslink DNA and proteins. Biotin-labeled DNA was detected using LightShift chemiluminescence EMSA kit (Pierce). The sequences of the probes are listed in Table S3.

### smFISH, phenotypic scoring, and statistical analysis

Single-molecule fluorescence in situ hybridization (smFISH) was performed as described previously (Topalidou et al., 2011). Imaging was conducted on a Zeiss Axio Observer Z1 inverted microscope with a CoolSNAP HQ2-FW camera (Photometrics, Tucson, AZ).

To examine the expression pattern of TRN markers, we grew animals at 20°C, examined them using the same microscope and recorded the percentages of TRN cells that express the fluorescent reporter in three independent experiments. The results are presented as aggregates; no significant differences were seen between replicates. For transgenic animals, at least three independent lines were examined.

Statistical significance was determined using the Student’s t-test for the majority of comparisons of two sets of data. For multiple comparisons, the Holm-Bonferroni method was used to correct the *p* values.

## Acknowledgement

We thank Alex Bounoutas for contributing to the initial studies on EGL-4 binding and Songtao Jia and Elizabeth Miller for sharing materials and reagents. We also thank Oliver Hobert, Richard Mann, and the members of our laboratory for helpful discussions and comments.

## Author Contributions

C.Z., F.Q.J., and B.L.T. performed experiments and analyzed the data. C.Z. and M.C. conceived the study and wrote the manuscript. J.W. performed the initial study about EGL-44 binding to TRN promoters. M.C. supervised the work and provided funding. F.Q.J. and B.L.T. contributed equally to the work.

